# Levels of Notch-regulated transcription are modulated by tissue movements at gastrulation

**DOI:** 10.1101/2021.09.15.460472

**Authors:** Julia Falo-Sanjuan, Sarah J. Bray

**Affiliations:** Department of Physiology, Development and Neuroscience, University of Cambridge, Downing Street, Cambridge CB2 3DY, UK

## Abstract

Cells sense and integrate external information from diverse sources that include mechanical cues. Shaping of tissues during development may thus require coordination between mechanical forces from morphogenesis and cell-cell signalling to confer appropriate changes in gene expression. By live-imaging Notch-induced transcription in real time we have discovered that morphogenetic movements during *Drosophila* gastrulation bring about an increase in activity-levels of a Notch responsive enhancer. Mutations that disrupt the timing of gastrulation resulted in concomitant delays in transcription up-regulation that correlated with the start of mesoderm invagination. As a similar gastrulation-induced effect was detected when transcription was elicited by the intracellular domain NICD, it cannot be attributed to forces exerted on Notch receptor activation. A Notch independent *vnd* enhancer also exhibited a modest gastrulation-induced activity increase in the same stripe of cells. Together, these observations argue that gastrulation-associated forces act on the nucleus to modulate transcription levels. This regulation was uncoupled when the complex linking the nucleoskeleton and cytoskeleton (LINC) was disrupted, indicating a likely conduit. We propose that the coupling between tissue level mechanics, arising from gastrulation, and enhancer activity represents a general mechanism for ensuring correct tissue specification during development and that Notch dependent enhancers are highly sensitive to this regulation.

## Introduction

Cells continuously sense and respond to their environment. This occurs via signalling pathways that detect and respond to external stimuli, such as morphogen gradients or direct cell-surface ligands. During development, information routed through these pathways feeds into networks of transcription factors that coordinate cell fates to pattern and regulate differentiation. At the same time, cells are exposed to mechanical forces from changes in tissue movements, deformations and stiffness. In order for tissues to develop and function correctly, there must be mechanisms that couple cell signaling with the mechanical environment. Indeed, there is evidence that changes in cell morphology can impact on gene expression ***(Alam et al. 2016; Guilluy et al. 2014)***, but this has been little explored in the context of developmental signaling.

Notch is a key developmental signalling pathway that transmits information between cells in contact. Prior to ligand binding, the ADAM10 cleavage site in Notch is hidden by the negative regulatory region (NRR). To reveal this site, ligand binding exerts a force on the receptor, leading to a displacement of the NRR ***(Gordon et al. 2007; Gordon et al. 2015)***. A second cleavage by γ-secretase then releases the intracellular domain, NICD, which forms a complex with a DNA-binding transcription factor and a co-activator to regulate transcription of target genes. In many developmental processes, Notch activation occurs contemporaneously with morphological changes which could modulate pathway activity so that signaling and tissue rearrangements are coordinated. For example, cell shape or tension changes in the neighbouring cells could impact on the forces exerted on the receptor to alter the amount of cleavage ***(Shaya et al. 2017)***. Alternatively, mechanical forces could alter the transcriptional output from Notch activation, by changing transport through nuclear pores or altering chromatin compaction ***(Boumendil et al. 2019; Gozalo et al. 2020)***.

To distinguish whether morphological events exert an influence on signaling, it is important to monitor the outputs in real-time when the changes are occurring. We have been investigating the onset of Notch activity in the *Drosophila* embryo using the MS2-MCP system to monitor the transcriptional response live. Notch activity initiates in a stripe of cells flanking the mesoderm, the mesectoderm (MSE), in the last cycle prior to gastrulation (nuclear cycle 14, nc14), and remains active throughout gastrulation. A notable feature of the transcriptional profiles from two Notch-responsive enhancers, *sim* and *m5/m8*, is that they undergo a transition approximately 50 min into nc14, when transcription levels increase approximately two-fold ***(Falo-Sanjuan et al. 2019)***. This transition occurs as the embryo undergoes gastrulation, making it plausible that mechanical forces or tissue reorganization at this stage are responsible for the increase in transcription.

Gastrulation initiates towards the end of nc14, approximately 3h post fertilization. During this process, the apical surface of the most ventral subset of mesoderm (ME) cells constricts and cells shorten along their apico-basal axis ***(Leptin and Grunewald 1990; Sweeton et al. 1991)***. This occurs in response to apical re-localization of Myosin II (MyoII), which is controlled by a GPCR (G-Protein Coupled Receptor) cascade ***(Sweeton et al. 1991; Manning et al. 2013; Parks and Wieschaus 1991; Barrett et al. 1997; Kolsch et al. 2007; Dawes-Hoang et al. 2005)***. As a consequence, the ‘ventral furrow’ is formed, which invaginates bringing the remainder of the mesoderm cells with it ***(Leptin and Grunewald 1990; Sweeton et al. 1991)***. Studies of the forces generated indicate that, although the force to invaginate the furrow is produced autonomously in the mesoderm by pulses of acto-myosin contractions ***(Martin et al. 2009)***, the mechanical properties of tissues adjacent to it, such as the MSE and neuroectoderm (NE), also change and may be important to allow invagination to occur ***(Rauzi et al. 2015)***.

We set out to investigate whether there is a causal relationship between morphological events occurring at gastrulation and the change in levels of Notch dependent transcription, using live imaging. Strikingly, we found a strong correlation between the start of mesoderm invagination and the time at which transcription levels increased. This change in transcription was delayed or absent when gastrulation was perturbed using different genetic mutations and manipulations. A Notch independent enhancer exhibited similar, albeit more modest effect, suggesting that the mechanism is a more general one. Futhermore, the Notch intracellular fragment, NICD, was also subject to similar regulation. The results indicate therefore that the mechanical context has a significant impact on the transcriptional outcome of Notch signaling but argue that this operates downstream of receptor activation and also affects other developmental enhancer(s). This type of coordination between tissue forces, developmental signalling and nuclear activity is thus likely to be of general importance for shaping tissues and organs as they form.

## Results

### Notch dependent transcription increases at gastrulation

During early *Drosophila* embryogenesis, Notch is active in the mesectoderm (MSE), a stripe of cells that border the mesoderm ***(Nambu et al. 1990; Morel and Schweisguth 2000; Morel et al. 2003)***. Activity commences midway through nc14 and continues during gastrulation, persisting in most MSE cells until they divide to form the precursors of the midline ***(Martín-Bermudo et al. 1995; Cowden and Levine 2002; Falo-Sanjuan et al. 2019)***. Measuring transcription live, using the MS2 system, reveals that there is a striking increase in the mean levels of transcription produced by Notch responsive enhancers during gastrulation. This increase occurs approximately 20 minutes after transcription initiates, which corresponds to 50 min after the start of nuclear cycle 14 (nc14). At this point, the transcription levels from two distinct Notch responsive mesectodermal enhancers, *m5/m8* and *sim* ***(Zinzen et al. 2006; Hong et al. 2013)***, almost double in magnitude (1**AB, *Falo-Sanjuan et al. 2019***). The change in magnitude occurs with different promoter combinations and with reporters inserted into different genomic positions (**Fig**. S1**AC****, *Falo-Sanjuan et al. 2019***), arguing it is a feature of the transcriptional response driven by the Notch responsive enhancers. However, there was no coordinated transition in levels when the profiles were aligned by the activity onset (**Fig**. S1**BD**), arguing that the increase is not dependent on the length of time that the enhancers have been active and thus may be related to a specific developmental event/time.

The observed increase in the mean transcriptional activity was not due to an increase in the number of active cells (**Fig**. 1**D**), but rather to a change in the profiles within individual nuclei. This differed in magnitude from nucleus to nucleus. Thus, when *m5/m8* transcription profiles from individual nuclei were analyzed, a clear, circa 2-fold, transition in levels was detected in 30-45% of the MSE nuclei (**Fig**. 1**C**). Others did not show such a major increase in levels, but several still manifest an inflection in the bursting at the equivalent time (**Fig**. 1**C**).

**Figure 1.**
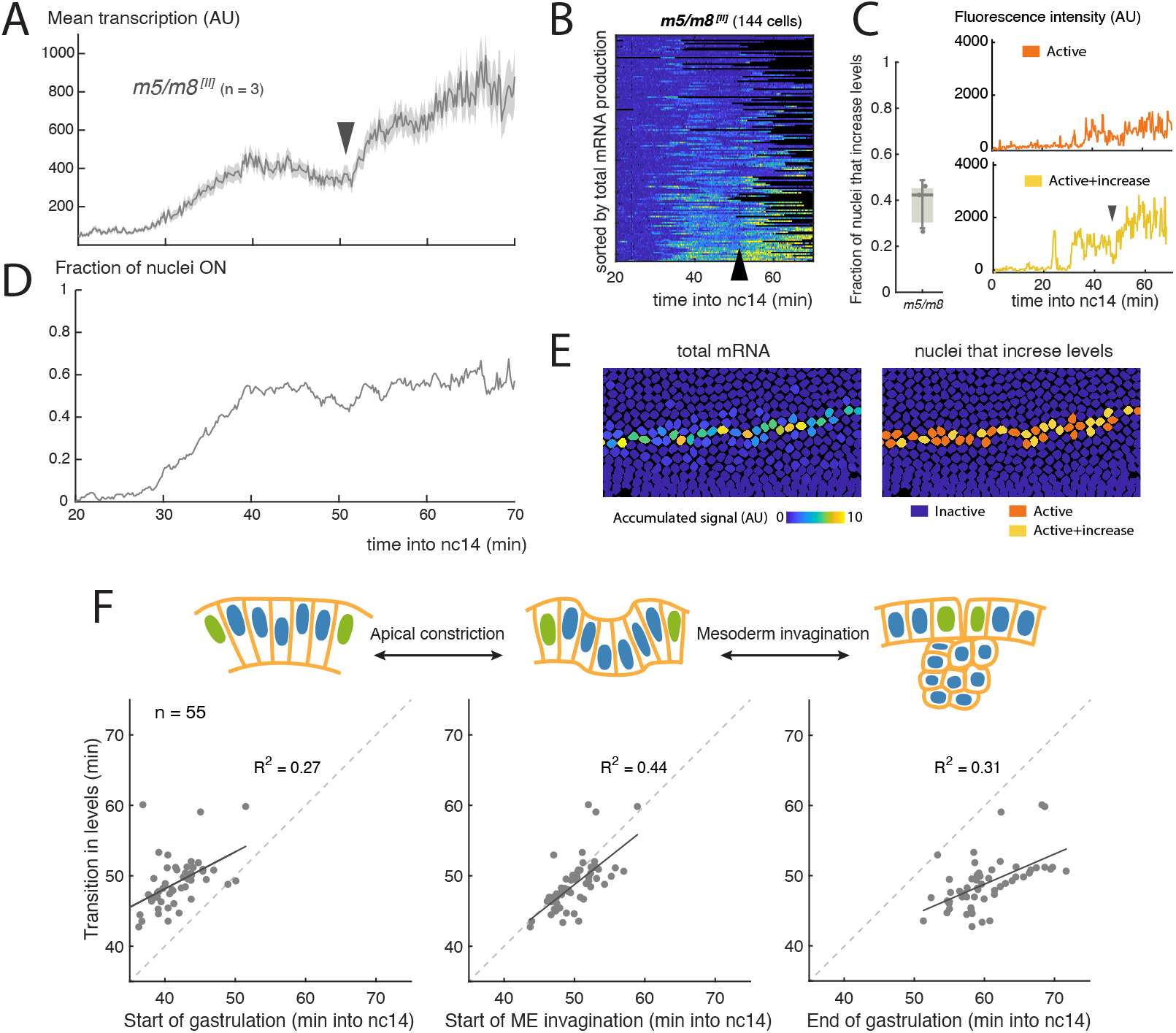
Increase in Notch-dependent transcription occurs at gastrulation. **A**) Mean profile of activity of the Notch responsive *m5/m8* ^*[II]*^ enhancer during nc14. **B**) Heatmap showing *m5/m8* ^*[II]*^ activity in all MSE cells over time, arrowhead indicates the transition point. **C**) Proportion of active cells in each embryo that increase levels of *m5/m8* ^*[II]*^ transcription at gastrulation (left; median, Q1/Q3 and SD shown) and examples of individual *m5/m8* ^*[II]*^ transcription traces with ∼2x increase (yellow) or inflection only (orange). **D**) Fraction of tracked nuclei that are actively transcribing during nc14. **E**) Still frame showing tracked nuclei color-coded for the total transcription produced by each nuclei (left) and by whether they exhibit ∼2x increase in levels (right). **F**) Correlation between the time at which mean levels of transcription increase in an embryo and the timing of 3 events during gastrulation. Each dot indicates an embryo from a collection of different Notch responsive enhancers, promoters and landing sites (n = 56 movies). All panels were obtained by re-analyzing data from ***Falo-Sanjuan et al. 2019***.

The transition in levels occurs whilst the embryo undergoes gastrulation (**Movie 1**), a process that starts midway through nc14 and lasts approximately 20 min. Gastrulation begins when the most ventral mesodermal cells constrict their apical surface to initiate ME invagination and results in the convergence of the two MSE stripes at the midline ***(Leptin and Grunewald 1990; Sweeton et al. 1991)***. Given the large scale morphogenetic movements involved, it is possible that these in some way influence the transcriptional activity. We therefore asked whether any of the changes that take place at gastrulation are correlated with the transition in transcription from the Notch responsive enhancers, by analyzing all the time-course data previously obtained from Notch responsive reporters (combining different enhancers, promoters and landing sites, n = 56 movies, ***Falo-Sanjuan et al. 2019***). On an embryo-by-embryo basis, we examined the relationship between the transcription transition-point and three different features during gastrulation: start of apical constriction, start of mesoderm invagination (defined as when mesectoderm cells first start to move ventrally) and the end of gastrulation (when all mesoderm cells have invaginated). Of the three, the highest correlation (both in terms of *R*^2^ and both events occurring at similar timepoints) was with the start of mesoderm invagination (**Fig**. 1**F**). Because of this correlation, one plausible model is that the morphogenetic movements from gastrulation are responsible for the increase in transcription levels.

### Events at gastrulation modulate Notch dependent transcription

To investigate whether morphological changes at gastrulation bring about the transition in Notch dependent transcription, we used a combination of approaches to disrupt gastrulation while live-imaging transcription from *m5/m8* ^*[II]*^, an insertion of the reporter on the second chromosome that responds robustly to Notch activity ***(Falo-Sanjuan et al. 2019)***. Gastrulation is coordinated by a signalling cascade that controls Myosin contractility, which in turn produces apical constriction of mesoderm cells to drive invagination (**Fig**. 2**A**) ***(Dawes-Hoang et al. 2005; Kolsch et al. 2007)***. First, we performed germline RNAi knockdowns (KD) to eliminate maternally-encoded proteins that act in the signaling cascade, namely α-Catenin (α-Cat), a key component of Adherens Junctions, which are required for apical RhoGEF2 and actomyosin relocalization ***(Dawes-Hoang et al. 2005; Kolsch et al. 2007)***, Concertina (Cta) and RhoGEF2, which are the Gα and GEF (Guanine nucleotide Exchange Factor) of the signaling cascade ***(Parks and Wieschaus 1991; Barrett et al. 1997)*** (**Fig**. 2**A**). The RNAi depletion produced morphological phenotypes consistent with those previously described, except in the case of RhoGEF2 ***(Parks and Wieschaus 1991; Barrett et al. 1997; Dawes-Hoang et al. 2005)***, where the depleted embryos were viable and lacked obvious morphological differences, suggesting a lower knock-down efficiency (**Fig**. S2**A**).

**Figure 2.**
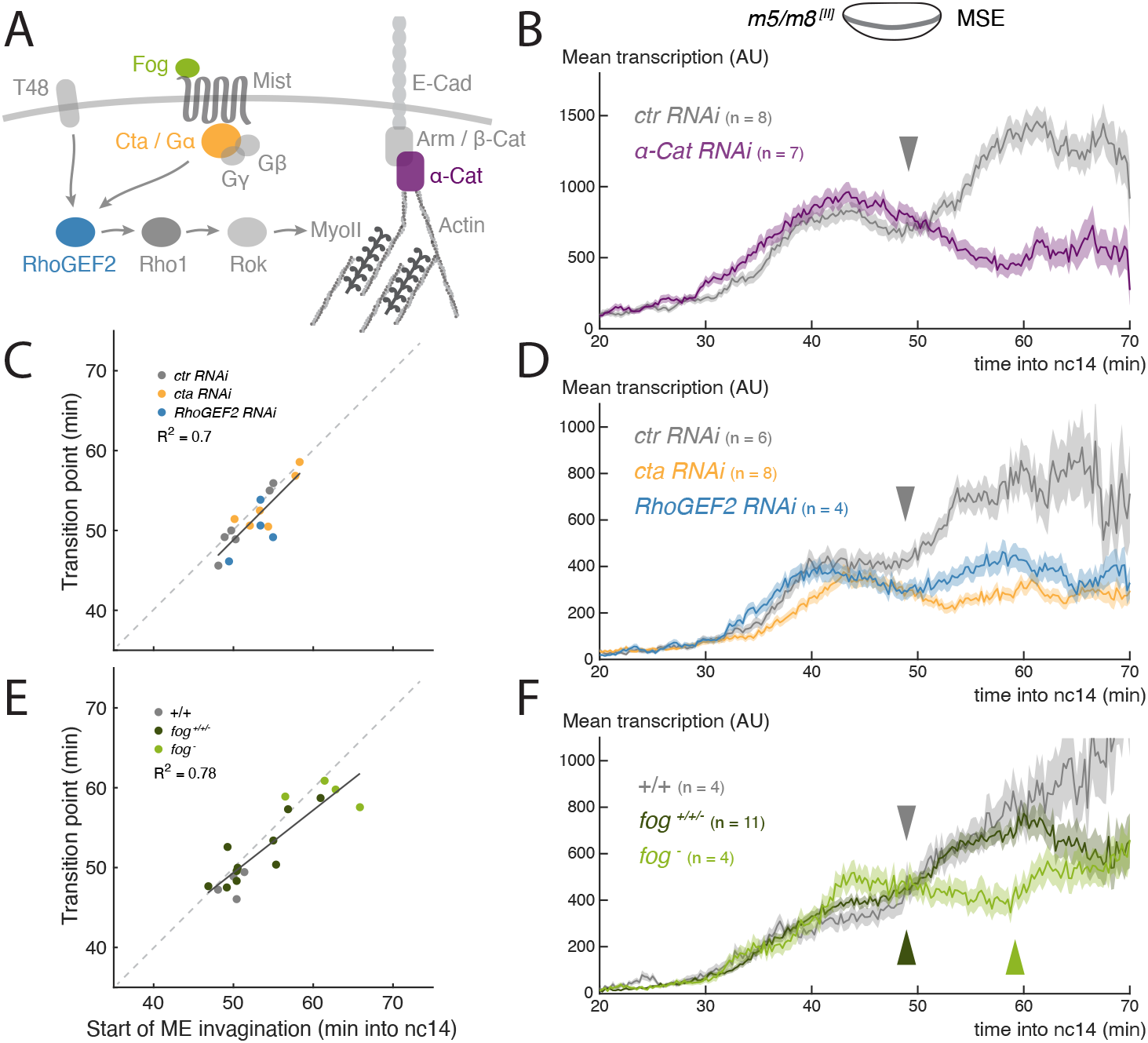
Disruption of gastrulation correlates with changes in the transition in Notch dependent transcription. **A**) Simplified scheme of the signalling cascade that controls MyoII contractility during *Drosophila* gastrulation. **B**) Mean profile of *m5/m8* ^*[II]*^ activity in *α-Cat* RNAi embryos compared to control embryos. **C**) Correlation between the start of invagination and transition in levels of transcription in each embryo, in *cta, RhoGEF2* and control RNAi embryos. **D**) Mean profile of *m5/m8* ^*[II]*^ activity in *cta, RhoGEF2* and control RNAi embryos. **E**) Correlation between the start of invagination and transition in levels of transcription in each embryo, in *fog* mutant embryos compared to control embryos and other non *fog* hemizygous embryos obtained from the same cross. **F**) Mean profile of *m5/m8* activity in *fog* mutant embryos compared to control embryos and other non *fog* hemizygous embryos obtained from the same cross. The transition in levels is delayed approximately 10 min in *fog* mutants (arrowheads). Mean transcription profiles show mean and SEM (shaded area) of MS2 fluorescent traces from all cells combined from multiple embryos (n embryo numbers indicated in each). *R*^2^ coefficients are calculated after pooling all points shown the same plot together. The transition point was only considered when a clear change in mean levels of transcription in an individual embryo could be observed, therefore the number of points in **C** and **E** could be smaller than the total number of embryos collected for each condition.

Of the three tested, α-Cat KD led to the most severe disruption of gastrulation, mesoderm cells failed to invaginate and divided externally (**Fig**. S2**ABC****, Movie 2**). Strikingly, no increase in *m5/m8* ^*[II]*^-directed transcription occurred in these embryos. Instead, the mean levels decreased after the initial activation, at the time when the increase in levels normally occurs, and then plateaued at a lower level (**Fig**. 2**B**, S3**B**). Likewise, the number of individual nuclei exhibiting an increase in levels was also greatly reduced (**Fig**. S3**A**). We note the mean levels of transcription from this reporter were not significantly reduced prior to gastrulation, unlike those from another reporter insertion and the endogenous genes that showed decreased levels also during cellularization ***(Falo-Sanjuan and Bray 2021)***. The difference likely arises because this *m5/m8* ^*[II]*^ insertion achieves maximal transcription at lower levels of Notch activity.

Depletion of either Cta or RhoGEF2 resulted in a slowing of gastrulation. Cta depletion also caused variable delays in the start of mesoderm invagination (**Fig**. S2**BC****, Movie 3**). In neither genotype was there a normal increase in transcription at the time of gastrulation. The mean levels remained similar to those prior to gastrulation and, on an individual nucleus basis, very few exhibited any marked change in levels (**Fig**. 2**D**, S3**CD**). Although there was no clear increase detected, the profiles retained an inflection point, when there was a transition between the early ‘peak’ levels and the later activity, manifest as a small dip in levels between the two phases. To assess whether this transition point was related to gastrulation events, the time when the transition occurred was plotted against the mile-stones for each embryo individually. The strongest correlation was with the start of mesoderm invagination (coefficient of *R*^2^ = 0.7, **Fig**. 2**C**), as it was for the increase in levels that occurs in wild-type embryos.

We next evaluated the consequences from mutations affecting the zygotically required gene *folded gastrulation* (*fog*), which encodes the ligand for the GPCR in the cascade regulating gastrulation ***(Costa et al. 1994; Dawes-Hoang et al. 2005)***. *fog*^*-*^ hemizygous embryos exhibited a delayed start of mesoderm invagination and slowed gastrulation overall (**Fig**. S2**BC****, Movie 2**). In these embryos, levels of transcription increased, although less than in normal embryos (**Fig**. 2**F**, S3**EF**). Notably, there was a significant 10 min delay in the time at which levels increased, from approximately 50 min to 60 min into nc14 (**Fig**. 2**F**). This time-point also correlated well (*R*^2^ = 0.78) with the start of mesoderm invagination when analyzed on an embryo-by embryo basis (**Fig**. 2**E**), similar to the transition-point in the RNAi depleted embryos.

Based on these genetic experiments, Notch-dependent transcription levels do not increase in conditions where gastrulation is blocked or slowed down. We hypothesize therefore that normal ‘fast’ gastrulation is required for the increase in levels and, since we observe a consistent correlation between the start of mesoderm invagination and the transition in transcription levels, it is likely that this is the causal step.

### Gastrulation also modulates Notch independent transcription

There are several different types of mechanisms that could explain a causal link between gastrulation and Notch-dependent transcription levels. As Notch activation involves a pulling force from Delta in adjacent cells ***(Gordon et al. 2015)***, one model is that increased cell surface tensions from the morphogenetic movements at gastrulation lead to increased Notch cleavage and NICD release. An alternative model is that the forces bring about a change in nuclear properties that more generally impact on transcription.

To distinguish these possibilities we took two approaches. First, we generated a transcriptional reporter using a Notch-independent enhancer from the *ventral nervous system defective* (*vnd*) gene. Unlike many of the other embryonic enhancers studied to date, the *vnd early embryonic enhancer* (*vndEEE*) is reported to be active throughout nc14 and to drive expression in a band of cells that overlap the MSE ***(Stathopoulos et al. 2002)***. As predicted, a new MS2 reporter containing this enhancer, referred to here as *vnd*, was active from the beginning of nc14 throughout gastrulation and recapitulated the spatial pattern of *vnd* (**Movie 4**). After a peak of transcription at the start of nc14, the mean levels of *vnd* transcription from all the active nuclei exhibited no increase in levels at gastrulation. Indeed, the overall mean decreased at this time. However, when the nuclei were separated according to their expression domain, an increase in mean levels was detectable specifically in MSE nuclei at around 50 min into nc14, similar to the *m5/m8* enhancer (**Fig**. 3**A**). No increase was detected in the other, NE domain (**Fig**. 3**B**). The proportion of nuclei that exhibited a clear increase in levels when considered individually was, however, considerably lower than for *m5/m8* (16% of MSE nuclei and 2% of NE nuclei, **Fig**. 3**C**). These results suggest that the activity of Notch independent as well as Notch dependent enhancers are affected at the time of gastrulation, albeit the effect is more modest for Notch independent activity, and that this property is limited to the mesectodermal cells.

**Figure 3.**
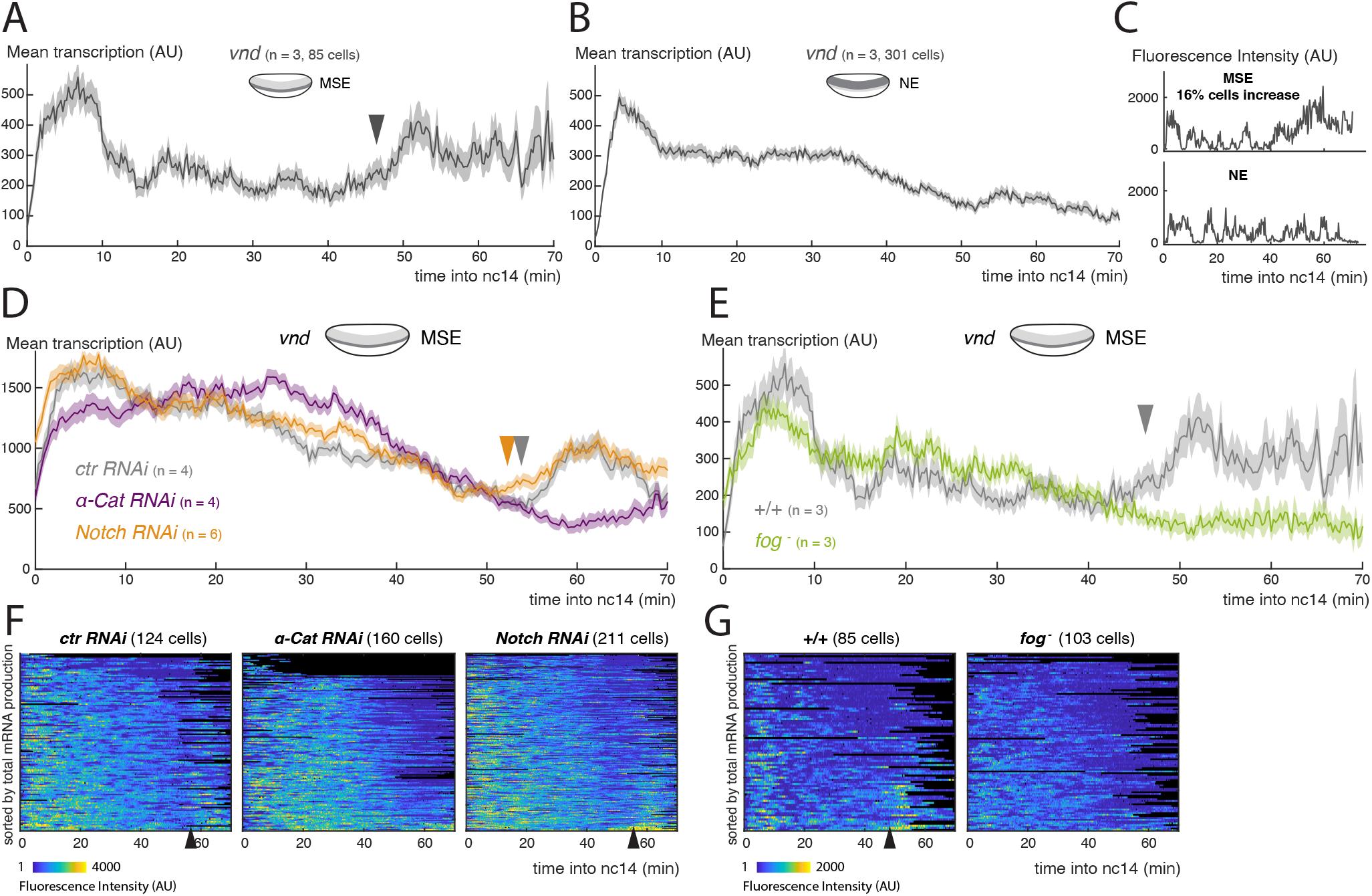
Activity of the Notch-independent *vnd* enhancer is modulated by gastrulation. **A**) Mean profile of transcription from *vnd* in MSE nuclei. **B**) Mean profile of transcription of *vnd* in NE nuclei. **C**) Examples of *vnd* transcription traces from MSE and NE, with (upper) and without (lower) a marked increase at the time of gastrulation. **D**) Mean profile of transcription from *vnd* in MSE nuclei in *α-Cat* and *Notch* RNAi compared to control embryos. **E**) Mean profile of transcription from *vnd* in MSE nuclei in *fog* mutant embryos compared to control embryos. **F**) Heatmaps showing *vnd* activity in all MSE cells over time in *α-Cat, Notch* and control RNAi embryos. **G**) Heatmaps showing *vnd* activity in all MSE cells over time in *fog*^*-*^ embryos compared to controls. Mean transcription profiles show mean and SEM (shaded area) of MS2 fluorescent traces from all cells combined from multiple embryos (n embryo numbers indicated in each). Arrowheads indicate increase in the mean levels of transcription.

We next investigated whether the increase in *vnd* activity in MSE nuclei was also linked to gastrulation, by measuring transcription in α-Cat depleted and *fog* mutant embryos. Neither genotype showed an increase in transcription in MSE cells (**Fig**. 3**D**-**G**), instead the levels remained at a plateau similar to that in the NE nuclei (**Fig**. 3**B**). Thus it appears that *vnd* transcription in MSE nuclei is also augmented due to gastrulation, in a similar manner to *m5/m8*-directed transcription. As there is no evidence that *vnd* is regulated by Notch ***(Markstein et al. 2004)***, this implies that gastrulation exerts effects on transcription independent of any effects on Notch activation. To verify that the modulation of *vnd* is not Notch dependent, we used an RNAi line to deplete Notch levels. As predicted, this treatment greatly reduced transcription from *m5/m8* ^*[II]*^ (**Fig**. S4**AB**). In contrast, there was no change in the timing of either the transition or the increase in levels from the *vnd* enhancer (**Fig**. 3**DF**). Together the results indicate that gastrulation also modulates the activity of a Notch independent *vnd* enhancer, suggesting that a general, rather than a Notch-specific, mechanism is involved.

### Regulation of Notch dependent transcription by gastrulation occurs downstream of pathway activation

The results with the *vnd* enhancer suggest that the gastrulation-induced changes in *m5/m8* transcription do not arise from increased Notch cleavage. As a second approach to investigate at which level of the pathway this modulation occurs, we examined whether gastrulation exerted any effects on transcription levels when the intracellular fragment NICD was expressed ectopically. To do so, we used an *eve2* transgene to direct expression of NICD in an orthogonal stripe overlapping the MSE, which is sufficient to drive ectopic transcription from *m5/m8* ***(Kosman and Small 1997; Cowden and Levine 2002; Falo-Sanjuan et al. 2019)***. The mean profile of *m5/m8* ^*[II]*^ transcription in MSE nuclei that were exposed to NICD had two peaks, the first shortly after the onset of NICD expression and the second during gastrulation. The transition between the two, the ‘trough’, slightly preceded the onset of mesoderm invagination. Thus the second ‘peak’ was initiated at the start of mesoderm invagination, characteristic of the gastrulation-induced increase in transcription in normal conditions. As MSE nuclei will be exposed to endogenous Notch signalling as well as the ectopic NICD, we also examined the profiles in the bordering NE nuclei. These exhibited a similar trough in levels prior to gastrulation, although the subsequent activity did not achieve the same levels as in the MSE nuclei (**Fig**. S5**B**). Together, these data support the model that gastrulation has an effect on *m5/m8* transcription that is independent of any influence on Notch cleavage.

To further verify that gastrulation-dependent changes occur independent from Notch cleavage, we monitored the transition in transcription levels elicited by NICD in embryos where the endogenous Notch was depleted by RNAi. No transcription was detected outside the domain of the *eve2* stripe (**Fig**. S5**A****, Movie 5**), confirming that Notch depletion was successful. Strikingly, the MSE nuclei within the stripe where NICD was expressed exhibited the same profile of activity in the *Notch RNAi* depleted embryos to those from control embryos with intact Notch. In particular, the levels of transcription increased at the time of gastrulation and to the same degree (**Fig**. 4**AC**). Thus, the transition in transcription levels at gastrulation occurs even in the absence of full length Notch, arguing the effects are downstream of receptor cleavage.

**Figure 4.**
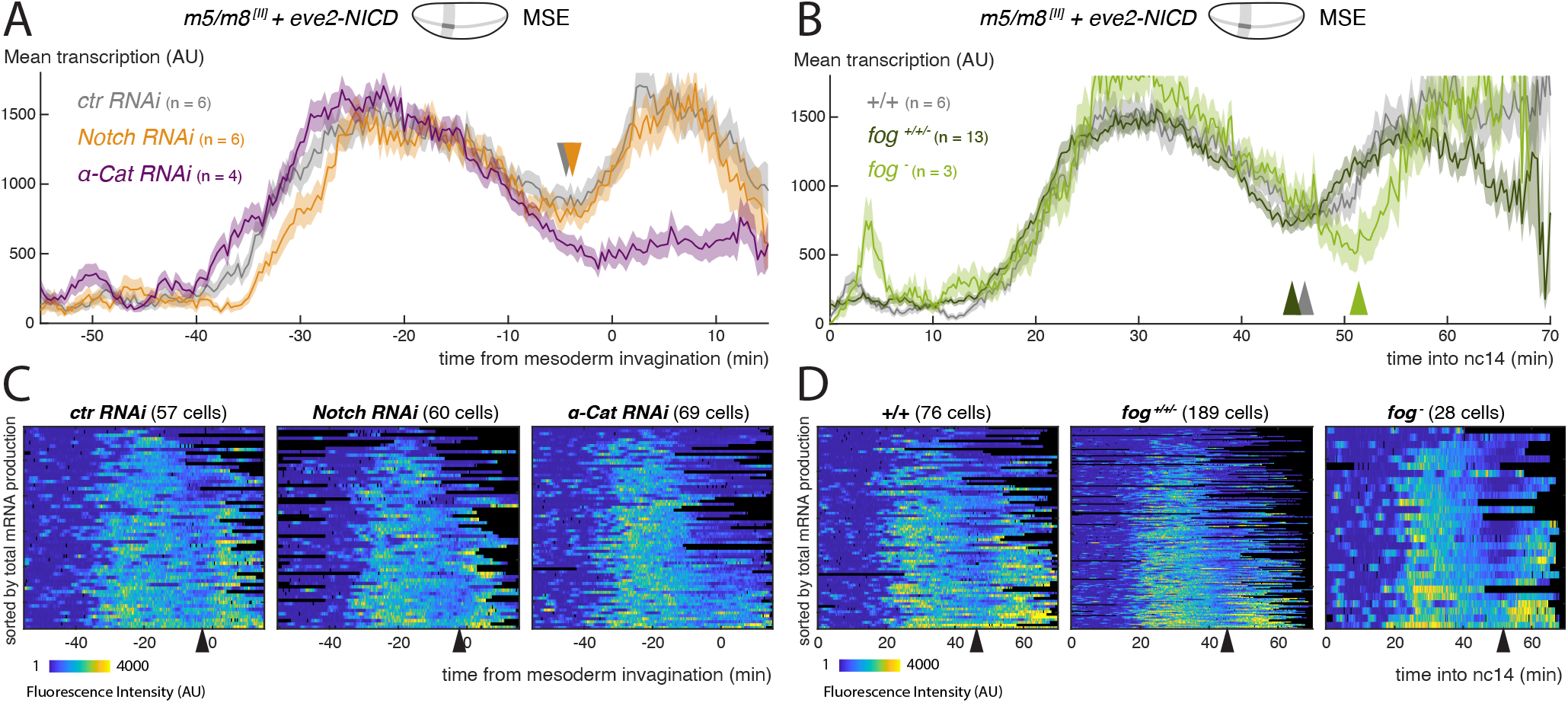
Modulation of Notch dependent transcription occurs downstream of pathway activation. **A**) Mean profile of transcription from *m5/m8* ^*[II]*^ in MSE nuclei, aligned by the time of ME invagination, in *α-Cat* and *Notch* RNAi compared to control embryos in the presence of ectopic NICD (produced by *eve2-NICD*). **B**) Mean profile of *m5/m8* ^*[II]*^ transcription in MSE nuclei with ectopic NICD in homozygous *fog* mutants compared to control or heterozygous embryos from the same cross (*fog*^*+/+/-*^). **C**) Heatmaps showing *m5/m8* ^*[II]*^ activity in MSE cells within *eve2-NICD* domain, aligned by the time of ME invagination, in *α-Cat, Notch* and control RNAi embryos. **D**) Heatmaps showing *m5/m8* ^*[II]*^ activity in MSE cells within *eve2-NICD* domain over time in *fog*^*-*^ embryos compared to control or heterozygous embryos obtained from the same cross (*fog*^*+/+/-*^). Mean transcription profiles show mean and SEM (shaded area) of MS2 fluorescent traces from all cells combined from multiple embryos (n embryo numbers indicated in each). Arrowheads indicate an increase in the mean levels of transcription.

To investigate whether the transition in transcription that occurs in the presence of NICD is directly related to gastrulation, we examined it in embryos where gastrulation was disrupted by α-Cat depletion or the *fog* allele, as above. In the α-Cat depleted embryos, where gastrulation was completely disrupted, there was no second ‘peak’ of transcription from *m5/m8* ^*[II]*^. Instead the levels remained at low levels from the time when mesoderm invagination would normally occur (**Fig**. 4**AC**). *fog* mutant embryos, in which gastrulation was delayed and slowed down, retained a second peak of *m5/m8* ^*[II]*^ transcription, but its onset was delayed in both MSE and NE nuclei (**Fig**. 4**BD**, S5**BC**). Notably, on an embryo by embryo basis, the delayed transition-point in the MSE correlated with the start of mesoderm invagination (*R*^2^ coefficient of 0.62) (**Fig**. S5**D**).

Altogether, the results suggest that the mechanics of gastrulation have an input into transcription in the MSE, that brings about an increase in the levels of mRNA produced. Because the effects on the *m5/m8* enhancer are more marked than those on *vnd*, Notch-dependent processes may be more sensitive. Furthermore, the consequences are strongest in MSE cells, suggesting that gastrulation exerts differential impacts across the ectoderm, which might be related to the gradients of forces and cell deformation that have been measured ***(Fuse et al. 2013; Rauzi et al. 2015)***.

### Disruption of LINC complex alters the profile of *m5/m8* transcription

Amongst mechanisms that could be responsible for the gastrulation-related increase in transcriptional activity in MSE nuclei, one hypothesis is that forces from cell rearrangements are transmitted through the cytoskeleton to the nucleus via the Linker of Nucleoskeleton and Cytoskeleton (LINC) complex (**Fig**. 5**A****, *Crisp et al. 2006***). We therefore set out to test the impact from disrupting components of the LINC complex on the transcription profile of *m5/m8* ^*[II]*^. Of the three conditions tested, only maternal knockdown of *klarsicht* (*klar*), which encodes a KASH protein, was successful, producing embryos with strongly reduced *klar* mRNA (**Fig**. S6**A**). Depletion of mRNA encoding SUN protein Klaroid (Koi) was unsuccessful and *Lamin C* mRNA knockdown produced no viable embryos. We therefore focused on the effects produced by *klar* mRNA depletion.

**Figure 5.**
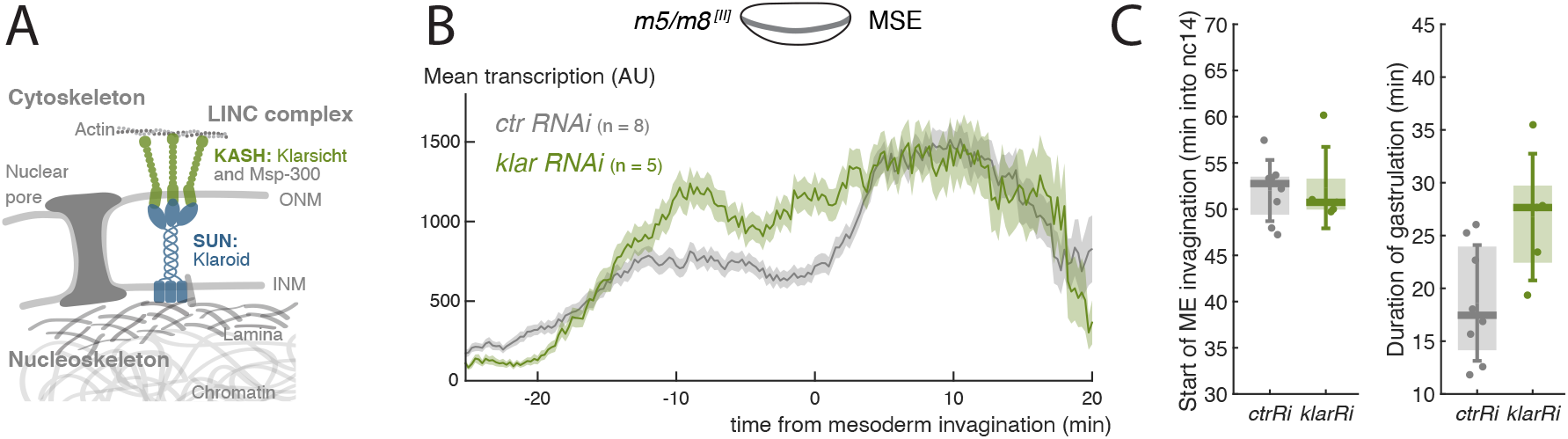
LINC complex disruption decouples gastrulation progression and changes in transcription. **A**) Scheme depicting LINC complex that connects cytoskeleton and nucleoskeleton, *Drosophila* KASH and SUN proteins Klarsicht and Klaroid are highlighted. **B**) Mean profile of transcription from *m5/m8* ^*[II]*^ in *klar* RNAi compared to control embryos. **C**) Onset of ME invagination (left) and duration of gastrulation in *klar* and control RNAi embryos. Boxplots show median, Q1/Q3 quartiles and SD. Control embryos are the same as shown in 2**B** and S2**B**.

If the LINC complex was involved in transcription modulation, we would expect its disruption to result in similar flat transcription levels to those seen when gastrulation is perturbed. Indeed, Klar disruption did yield flatter levels of *m5/m8* ^*[II]*^ transcription, but this occurred in the opposite way than predicted, because the early phase of transcription was slightly elevated and there was no subsequent transition or reduction in the late phase (**Fig**. 5**B**, S6**B**). As there was no distinct transition at the time of mesoderm invagination, this could indicate an uncoupling of the transcription profiles from the gastrulation movements caused by LINC complex disruption. However, these embryos exhibited longer and less robust gastrulation (**Fig**. 5**C**, S6**C**). As other manipulations that slow gastrulation appeared to have a dampening effect on the transition in transcription levels (see earlier), it is possible that the alterations in gastrulation could explain the lack of any increase in transcription, rather than a direct role of Klar in this process. Nevertheless, the change in profile from the *klar* mRNA knock-down suggest that the LINC complex helps couple the gastrulation movements with transcription.

## Discussion

Morphogenetic processes integrate with programs of transcriptional regulation during animal development, but little is known about crosstalk between mechanics and transcription, particularly in a native *in vivo* context. Here we have investigated a role of gastrulation in modulating transcription levels in *Drosophila* embryos. Using the MS2 system to quantitatively image transcription live, a clear transition in the mean levels of transcription from the Notch responsive *m5/m8* enhancer could be detected, and to a lesser degree from the Twist and Dorsal responsive *vnd* enhancer. This transition correlated with the start of mesoderm invagination and was delayed or absent in embryos where gastrulation was perturbed by different manipulations. In conditions where there was a delay, such as in *fog* mutants, the transition in levels was correlated with a delay in mesoderm invagination.

A plausible link between mechanics and Notch activity would be that force is required for Notch cleavage ***(Gordon et al. 2015)***. By increasing membrane tension or by rearranging cell contacts, the morphogenetic movements associated with gastrulation could enhance the levels of NICD released. However, two observations make this scenario unlikely as the primary cause for the effects on transcription at mesoderm invagination. First, the same effect, but to a smaller degree, was also observed with the *vnd* enhancer, which exhibited an increase in levels at the time of gastrulation that was correspondingly perturbed in *fog* mutants or α-Cat depleted embryos. Second, embryos in which full length Notch was maternally depleted while ectopically expressing NICD, which bypasses the need for force-mediated activation, exhibited similar gastrulation-dependent transition in levels. If the initial decline in levels in the presence of NICD involves an intrinsic attenuation mechanism, as has recently been proposed ***(Viswanathan et al. 2021)***, this must then be overcome in a gastrulation dependent manner. Altogether, our results argue in favour of a model where the cell movements associated with gastrulation have a direct impact on transcription in the nucleus.

A common feature of both *m5/m8* and *vnd* enhancers is that their activity in MSE cells is more profoundly influenced by gastrulation than elsewhere in the embryo. This could be due to differences in the context of transcription factors and other regulators present in the MSE cells, that make them more sensitive, or to variations in the magnitude of the mechanical forces exerted by gastrulation. In favor of the latter, ectoderm cells are thought to have different mechanical properties to the mesoderm and lateral cells, which contribute to the normal progression of gastrulation ***(Rauzi et al. 2015)***. We therefore hypothesize that transcription levels can be modulated by the forces exerted on cells, and that this effect may be specific to certain classes of enhancers. Here we have found that although both *m5/m8* and *vnd* enhancers show characteristics of such regulation, *m5/m8* appears the more sensitive. We propose that mechanisms acting at the level of the nucleus will be responsible for transmitting this force-mediated regulation.

To exert effects on nuclear functions, the mechanical changes induced by gastrulation need to be transmitted to the nucleus. The mechanotransduction cascade is thought to involve the cytoskeleton and the LINC complex ***(Chang et al. 2015)***. Along with myosin, the LINC complex is proposed to transmit information about substrate stiffness to the transcriptional machinery ***(Alam et al. 2016)***. A similar mechanism could be involved in the gastrulating embryo to transduce mechanical information at the cell level to changes in transcription, based on the fact that coupling between transcription and gastrulation was no longer evident when the LINC complex was depleted. Although the fact that there are also changes in gastrulation under these conditions make it difficult to conclude that there is a direct effect of the LINC complex, the results are nevertheless consistent with the model.

There is increasing evidence that substrate stiffness and mechanical forces can influence cell fate decisions **(Roy2018; *Alam et al. 2016; Wei et al. 2015; Muncie et al. 2020)***. These data showing a connection between gastrulation and transcription levels support the suggestion that there are mechanisms connecting the mechanical properties experienced by cells with the transcriptional output. Such a connectivity is likely to be of major significance in building tissues and organs, to ensure that the correct structure and shape are adopted in the context of the developing organism.

## Methods

### Fly strains and genetics

The MS2 reporter *m5/m8* ^*[II]*^ (*m5/m8-peve-24xMS2-lacZ-SV40*[25C]), inserted in the landing site attP40-25C in the second chromosome ***(Bischof et al. 2013)***, other reporters used in **Fig**. 1 and the *eve2-FRT-STOP-FRT-NICD*[51D] construct used to ectopically express NICD have been previously described ***(Falo-Sanjuan et al. 2019)***. *vnd-MS2* (*vndEEE-peve-24xMS2-lacZ-SV40*) was generated by replacing *m5/m8* for the *vndEEE* enhancer, as defined by ***Stathopoulos et al. 2002***, using primers *GGGAAGCTTGGGTAAGCACAAGGATTCC* and *GGGACCGGTCGAATAAGCTG-CAAGGAGATC* with *Hind*III and *Age*I sites. The resulting plasmid was inserted in the same attP landing site (attP40-25C) as *m5/m8* by φC31 mediated recombination ***(Bischof et al. 2013)***. Full genotypes of used lines are detailed in **Table** 1.

**Table 1.** Full genotypes and sources of used *Drosophila* lines.

| Name (Chr) | Full genotype | Source |
| --- | --- | --- |
| *His2Av::RFP* (III) | w[*];P{w[+mC]=His2Av-mRFP1}III.1 | BDSC #23650 |
| *His2Av::RFP; nos-MCP::GFP* | y[1] w[*]; P{w[+mC]=His2Av-mRFP1}II.2; P{w[+mC]=nos-MCP.EGFP}2 | BDSC #60340 |
| αTub-Gal4::VP16 (II) | w[*]; P{w[+mC]=matalpha4-GAL-VP16}V2H | BDSC #7062 |
| *ovo-FLP* (I) | P{w[+mC]=ovo-FLP.R}M1A, w[*] | BDSC #8727 |
| *betaTub85D-FLP* (I) | P{ry[+t7.2]=betaTub85D-FLP}1; ry[506] | BDSC #7196 |
| *m5/m8-MS2* (II) | w; P{w[+mC]=m5/m8-peve-24xMS2-lacZ-SV40}attP40 | (*Falo-Sanjuan et al. 2019*) |
| *vnd-MS2* (II) | w; P{w[+mC]=vndEEE-peve-24xMS2-lacZ-SV40}attP40 | This study |
| *eve2-NICD* (II) | w; P{w[+mC]=2xeve2*-FRT-STOP-FRT-NICD-eve3'UTR}attP51D | (*Falo-Sanjuan et al. 2019*) |
| *fog[S4]* (I) | y[1] fog[S4]/FM7c | BDSC #2100 |
| *w* RNAi Valium22 (III) | y[1] sc[*] v[1]; P{y[+t7.7] v[+t1.8]=TRiP.GL00094}attP2 | BDSC #35573 |
| α-Cat RNAi Valium20 (III) | y[1] sc[*] v[1] sev[21]; P{y[+t7.7] v[+t1.8]=TRiP.HMS00317}attP2 | BDSC #33430 |
| *cta* RNAi Valium20 (II) | y[1] sc[*] v[1]; P{y[+t7.7] v[+t1.8]=TRiP.HMC03421}attP40 | BDSC #51848 |
| *RhoGEF2* RNAi Valium20 (III) | y[1] sc[*] v[1]; P{y[+t7.7] v[+t1.8]=TRiP.HMS01118}attP2 | BDSC #34643 |
| *Notch* RNAi Valium20 (III) | y[1] v[1]; P{y[+t7.7] v[+t1.8]=TRiP.HMS00009}attP2 | BDSC #33616 |
| *klar* RNAi Valium20 (III) | y[1] sc[*] v[1] sev[21]; P{y[+t7.7] v[+t1.8]=TRiP.HMS01612}attP2 | BDSC #36721 |
| *koi* RNAi Valium20 (II) | y[1] sc[*] v[1] sev[21]; P{y[+t7.7] v[+t1.8]=TRiP.HMS02172}attP40 | BDSC #40924 |
| *lamC* RNAi Valium20 (II) | y[1] sc[*] v[1] sev[21]; P{y[+t7.7] v[+t1.8]=TRiP.HMC04816}attP40 | BDSC #57501 |

Crosses performed for each experiment are detailed in **Table** 2.

**Table 2.** Crosses to obtain embryos used in each experiment.

| Cross | Figure |
| --- | --- |
| ♀ αTub-Gal4::VP16 / + ; His2Av::RFP, nos-MCP::GFP / UASp-w RNAi<br>♂ m5/m8-peve-MS2-lacZ-SV40[attP40,II] | 2, S2, S4, 5, S6 |
| ♀ αTub-Gal4::VP16 / + ; His2Av::RFP, nos-MCP::GFP / UASp-α-Cat RNAi<br>♂ m5/m8-peve-MS2-lacZ-SV40[attP40,II] | 2, S2 |
| ♀ His2Av::RFP, nos-MCP::GFP (III) x ♂ m5/m8-peve-MS2-lacZ-SV40[attP40,II] | 2, S2 |
| ♀ fog[S4] / FM6 ;; His2Av::RFP, nos-MCP::GFP x ♂ m5/m8-peve-MS2-lacZ-SV40[attP2 40,II] | 2, S2 |
| ♀ αTub-Gal4::VP16 / UASp-cta RNAi ; His2Av::RFP, nos-MCP::GFP / +<br>♂ m5/m8-peve-MS2-lacZ-SV40[attP40,II] | 2, S2 |
| ♀ αTub-Gal4::VP16 / + ; His2Av::RFP, nos-MCP::GFP / UASp-RhoGEF2 RNAi<br>♂ m5/m8-peve-MS2-lacZ-SV40[attP40,II] | 2, S2 |
| ♀ His2Av::RFP, nos-MCP::GFP (III) x ♂ vndEEE-peve-MS2-lacZ-SV40[attP40,II] | 3, S4 |
| ♀ αTub-Gal4::VP16 / + ; His2Av::RFP, nos-MCP::GFP / UASp-w RNAi<br>♂ vndEEE-peve-MS2-lacZ-SV40[attP40,II] | 3, S4 |
| ♀ αTub-Gal4::VP16 / + ; His2Av::RFP, nos-MCP::GFP / UASp-α-Cat RNAi<br>♂ vndEEE-peve-MS2-lacZ-SV40[attP40,II] | 3, S4 |
| ♀ αTub-Gal4::VP16 / + ; His2Av::RFP, nos-MCP::GFP / UASp-Notch RNAi<br>♂ m5/m8-peve-MS2-lacZ-SV40[attP40,II] | S4 |
| ♀ αTub-Gal4::VP16 / + ; His2Av::RFP, nos-MCP::GFP / UASp-Notch RNAi<br>♂ vndEEE-peve-MS2-lacZ-SV40[attP40,II] | 3, S4 |
| ♀ fog[S4] / FM6 ;; His2Av::RFP, nos-MCP::GFP x ♂ vndEEE-peve-MS2-lacZ-SV40[attP40,II] | 3, S4 |
| ♀ ovo-FLP / + ; eve2-FRT-STOP-FRT-NICD / + ; His2Av::RFP, nos-MCP::GFP / +<br>♂ m5/m8-peve-MS2-lacZ-SV40[attP40,II] | 4, S5 |
| ♀ ovo-FLP / fog[S4] ; eve2-FRT-STOP-FRT-NICD / + ; His2Av::RFP, nos-MCP::GFP / +<br>♂ m5/m8-peve-MS2-lacZ-SV40[attP40,II] | 4, S5 |
| ♀ αTub-Gal4::VP16 / + ; His2Av::RFP, nos-MCP::GFP / UASp-w RNAi<br>♂ βTub85D-FLP ; m5/m8-peve-MS2-lacZ-SV40, eve2-FRT-STOP-FRT-NICD / + | 4, S5 |
| ♀ αTub-Gal4::VP16 / + ; His2Av::RFP, nos-MCP::GFP / UASp-Notch RNAi<br>♂ βTub85D-FLP ; m5/m8-peve-MS2-lacZ-SV40, eve2-FRT-STOP-FRT-NICD / + | 4, S5 |
| ♀ αTub-Gal4::VP16 / + ; His2Av::RFP, nos-MCP::GFP / UASp-α-Cat RNAi<br>♂ βTub85D-FLP ; m5/m8-peve-MS2-lacZ-SV40, eve2-FRT-STOP-FRT-NICD / + | 4, S5 |
| ♀ x ♂ αTub-Gal4::VP16 / + ; UASp-w RNAi / + | S6 |
| ♀ x ♂ αTub-Gal4::VP16 / UASp-koi RNAi ; + / + | S6 |
| ♀ x ♂ αTub-Gal4::VP16 / + ; UASp-klar RNAi / + | S6 |
| ♀ αTub-Gal4::VP16 / + ; His2Av::RFP, nos-MCP::GFP / UASp-klar RNAi<br>♂ m5/m8-peve-MS2-lacZ-SV40[attP40,II] | 5 |

#### fog mutant background

To test expression from *m5/m8* and *vnd* in the *fog* mutant background, third chromosome recombinants *His2av-RFP, nos-MCP-GFP* ***(Falo-Sanjuan et al. 2019)*** were combined with a *fog* null allele (*fog[S4]*, BDSC #2100). Control embryos were obtained by crossing *His2av-RFP, nos-MCP-GFP* females with *m5/m8* or *vnd* males. Hemizygous *fog* mutant embryos were obtained by crossing *fog[S4]* / *FM6* ;; *His2av-RFP, nos-MCP-GFP* with *m5/m8* or *vnd* males and they were recognized by the presence of ectopic folds after gastrulation and embryonic lethality. Other embryos obtained from the same cross were grouped together and labelled *fog* ^*+/+/-*^. To combine the *fog* background with ectopic NICD produced by *eve2-NICD, fog[S4]* was combined with *eve2-FRT-STOP-FRT-NICD* and crossed with *ovo-FLP* ;; *His2av-RFP, nos-MCP-GFP*. F1 females containing all transgenes were crossed with *m5/m8-MS2* males; and induce removal of the FRT-STOP-FRT cassete in the germline so that NICD is produced in the *eve2* pattern in F2 embryos. Controls for this experiment were obtained by crossing *eve2-FRT-STOP-FRT-NICD* with *ovo-FLP* ;; *His2av-RFP, nos-MCP-GFP* and F1 females with *m5/m8-MS2* males.

#### Maternal KD

The maternal driver *αTub-Gal4::VP16* (BDSC # 7062) was combined with *His2av-RFP, nos-MCP-GFP* to generate *αTub-Gal4::VP16* ; *His2Av-RFP, nos-MCP-GFP*. To knock down genes from the maternal germline, this was crossed with with UASp-RNAi lines detailed in **Table** 1 or *UASp-w-RNAi* as control (BDSC #35573). Females *αTub-Gal4::VP16* / + ; *His2Av-RFP, nos-MCP-GFP* / *UASp-RNAi* or *αTub-Gal4::VP16* / *UASp-RNAi* ; *His2Av-RFP, nos-MCP-GFP* / + were crossed with *m5/m8-MS2* or *vnd-MS2* to obtain the experimental embryos. To combine this with ectopic NICD, *m5/m8, eve2-FRT-STOP-FRT-NICD* recombinants ***(Falo-Sanjuan et al. 2019)*** had been previously crossed with *βTub-FLP* and males *βTub-FLP* / Y ; *m5/m8,eve2-FRT-STOP-FRT-NICD* / +, which induce FRT-STOP-FRT removal in the germline, crossed with females in which germline KD occurs. In this way, the resulting embryos express both MS2 and *eve2-NICD*, and candidate genes have been maternally knocked down.

To quantify the degree of maternal Klar and Koi KD, *αTub-Gal4::VP16* was crossed with the same lines and F2 embryos were collected for RT-qPCR.

### mRNA extraction and qPCR

RT-qPCR quantification of maternal KD was performed as previously described ***(Falo-Sanjuan and Bray 2021)***. Embryos were dechorionated in bleach and early embryos (pre-nc10) / eggs were selected in Voltalef medium. Pools of 15-20 embryos of each genotype were transfered to eppendorf tubes and dissociated in TRI Reagent (Sigma) with a plastic pestle. mRNA was extracted by adding chloroform, 10 min centrifugation at 4C and let to precipitate with isopropanol overnight. DNA was then pelleted by 10 min centrifugation at 4C, washed in 70% ethanol, dried and resuspended in DEPC-treated water. Approximately 2 μg of RNA from each sample were DNAse treated with the DNA-free™ DNA Removal Kit (Invitrogen) in the presence of RiboLock RNase Inhibitor (Thermo Scientific). 1 μg of DNA-free RNA was then used for reverse transcription using M-MLV Reverse Transcriptase (Promega) in the presence of RiboLock. RT-qPCR was performed using SYBR Green Mastermix (Sigma) and primers detailed in **Table** 3.

**Table 3.**
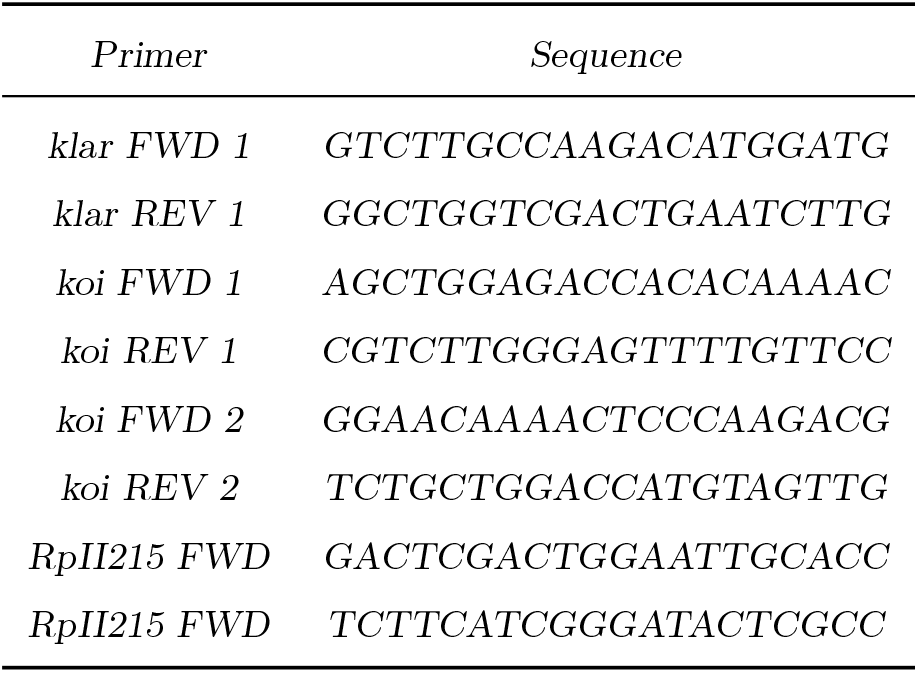
Primers used for qPCR.

### Live imaging

Embryos were collected on apple juice agar plates with yeast paste, dechorionated in bleach and mounted in Voltalef medium (Samaro) between a semi-permeable membrane and a coverslip. The ventral side of the embryo was facing the coverslip. Movies were acquired in a Leica SP8 confocal using a 40x apochromatic 1.3 objective, zoom x2 and 400×400px size (providing an XY resolution of 0.36 μm/px), 12 bit depth, 400 Hz image acquisition frequency and pinhole of 4 airy units. 27, 29 or 32 × 1 μm slices were collected, with total acquisition time of 15-20s per frame, depending on the experiment. Different genetic conditions are only compared in the same plot if the same microscope and settings were used.

### Image analysis

#### Tracking nuclei and MS2 quantification

Movies were analyzed using custom MATLAB (MATLAB R2020a, MathWorks) scripts that have been previously described ***(Falo-Sanjuan et al. 2019; Falo-Sanjuan and Bray 2021)***.

Briefly, the His2Av-RFP signal was used to segment and track the nuclei in 3D. Each 3D stack was first filtered using a 3D median filter of 3 and increasing the contrast based on the intensity profile of each frame to account for bleaching. This was followed by Fourier transform log filter to enhance round objects ***(Garcia et al. 2013)***, and segmentation by applying a fixed intensity threshold, which was empirically determined. Filters to fill holes in objects, exclusion based on size and 3D watershed accounting for anisotropic voxel sizes ***(Mishchenko 2015)*** were used to remove miss-segmented nuclei and split merged nuclei. The final segmented stack was obtained by filtering by size again and and thickening each object. Lastly, each nuclei in the segmented stack was labelled and the position of each centroid (in X, Y and Z) was calculated for tracking.

Nuclei were tracked in 3D by finding the nearest object (minimum Euclidean distance between two centroids in space) in the previous 2 frames which was closer than 6 μm. If no object was found, that nucleus was kept with a new label. If more than one object was ‘matched’ with the same one in one of the previous 2 frames, both were kept with new labels.

After tracking, the positions of all pixels from each nucleus in each frame were used to measure the maximum fluorescence value in the GFP channel, which was used as a proxy of the spot fluorescence. Note that when a spot cannot be detected by eye this method detects only background, but the signal:background ratio is high enough that the subsequent analysis allows to classify confidently when the maximum value is really representing a spot.

### Quantifying gastrulation progression and changes in transcription levels

Start of apical constriction, start of mesoderm invagination and end of gastrulation were manually defined for each embryo from plots showing the movement of MSE cells. Transcribing nuclei in each region were selected and the average movement of their centroids in the Y axis, corresponding to the DV axis in the embryo, was calculated. This produces a plot with one or two peaks of movement. A large peak of movement is produced during ME invagination, as MSE cells move ventrally. If the embryo is mounted completely ventrally, only this peak of movement is observed. If the orientation of the embryo is tilted, the whole embryo rolls inside the vitelline membrane during apical constriction, which is detected as an earlier peak of movement of cells dorsally. After that, the second peak of movement corresponds with mesoderm invagination. The transition between the two movements, corresponds to and was used to define the start of ME invagination. Similarly, transition point in mean transcription levels was manually defined from each plot showing the mean fluorescence of selected cells.

### Data processing and statistical analysis

#### MS2 data processing

Processing of MS2 data (definition of active nuclei and normalization for bleaching) has been carried out as described in our previous work ***(Falo-Sanjuan et al. 2019)***. After the fluorescent trace of each nucleus over time was obtained, only nuclei tracked for more than 10 frames were retained. Then, nuclei were classified as inactive or active. To do so, the average of all nuclei (active and inactive) was calculated over time and fitted to a straight line. A median filter of 3 was applied to each nucleus over time to smooth the trace and ON periods were considered when fluorescent values were 1.2 times the baseline at each time point. This produced an initial segregation of active (nuclei ON for at least 5 frames) and inactive nuclei. These parameters were determined empirically on the basis that the filters retained nuclei with spots close to background levels and excluded false positives from bright background pixels. The mean fluorescence from MCP-GFP in the inactive nuclei was then used to define the background baseline and active nuclei were segregated again in the same manner. The final fluorescence values in the active nuclei were calculated by removing the fitted baseline from the maximum intensity value for each, and normalizing for the percentage that the MCP-GFP fluorescence in inactive nuclei decreased over time to account for the loss of fluorescence due to bleaching. Nuclei active in cycles before nc14 were discarded based on the timing of their activation.

In all movies, time into nc14 was considered from the end of the 13th syncytial division. To avoid early stochastic activity, transcription was considered from 15 min into nc14 in most experiments, except for expression in the presence of maternal Gal4, in which it was considered from 30 min to exclude earlier stochastic activity induced by Gal4VP16. For *vnd*, transcription was considered all throughout nc14. The total mRNA output (in AU) was obtained by adding all the normalized transcription values for each cell in a defined time period. In plots were cells were classified as ‘active’ or ‘active+increasing’, nuclei which increase in levels were defined if the average intensity from 15 min of the start of ME invagination was higher than the average of all active nuclei at that point and at least 1.7 times higher than the average intensity in the 15 min prior to ME invagination. Using these values, most nuclei were classified in a way that matched what could be observed by looking directly at the transcription traces.

#### Statistical analysis

In figures and figure legends, n number indicates number of embryos imaged for each biological condition. Where appropriate, n number next to heatmaps indicates total number of cells combining all embryos for each biological condition. Plots showing mean levels of transcription and SEM (standard error of the mean) combine all traces from multiple embryos from the same biological condition.

### Reagents and software availability

The MATLAB app to track nuclei, quantify MS2 traces and define properties of gastrulation can be obtained at https://github.com/BrayLab/LiveTrx.

## Movies

**Movie 1 - Activity of *m5/m8* during gastrulation**. Movie showing His2Av-RFP channel (orthogonal view, left ; and maximum intensity projection, center) and MCP-GFP channel with transcription directed by *m5/m8* ^*[II]*^ (maximum intensity projection, right) in control embryos. The top row shows orthogonal view in the His2Av-RFP channel and maximum Y projection in the MCP-GFP channel. 0.36 μm/px XY resolution, 29×1μm slices and time resolution of 15s/frame. Anterior to the left; embryo imaged from the ventral side. Time indicates minutes from the beginning of nc14. Bottom plot shows mean transcription produced by *m5/m8* ^*[II]*^ in this embryo, synchronized with the movie to show the increase in activity occurs as the embryo undergoes gastrulation.

**Movie 2 - Effects on gastrulation of *α-Cat RNAi* and *fog* mutant background**. Movies showing His2Av-RFP channel (orthogonal view, left ; and maximum intensity projection, center) and MCP-GFP channel with transcription directed by *m5/m8* ^*[II]*^ (maximum intensity projection, right) in *α-Cat RNAi* (top) and *fog* ^*-*^ (bottom) embryos. 0.36 μm/px XY resolution, 27×1μm (*α-Cat RNAi*) and 29×1μm (*fog* ^*-*^) slices and time resolution of 15s/frame. Anterior to the left; embryo imaged from the ventral side. Time indicates minutes from the beginning of nc14.

**Movie 3 - Effects on gastrulation of *cta* and *RhoGEF2 RNAi***. Movies showing His2Av-RFP channel (orthogonal view, left ; and maximum intensity projection, center) and MCP-GFP channel with transcription directed by *m5/m8* ^*[II]*^ (maximum intensity projection, right) in *cta* (top) and *RhoGEF2 RNAi* (bottom) embryos. 0.36 μm/px XY resolution, 29×1μm slices and time resolution of 15s/frame. Anterior to the left; embryo imaged from the ventral side. Time indicates minutes from the beginning of nc14.

**Movie 4 - Expression of *vnd* in control embryos**. Movies showing His2Av-RFP channel (orthogonal view, left ; and maximum intensity projection, center) and MCP-GFP channel with transcription directed by *vnd* (maximum intensity projection, right) in control embryos. 0.36 μm/px XY resolution, 29×1μm slices and time resolution of 15s/frame. Anterior to the left; embryo imaged from the ventral side. Time indicates minutes from the beginning of nc14.

**Movie 5 - Expression of *m5/m8*** ^***[II]***^ **upon *eve2-NICD* expression in control, *Notch* and *α-Cat RNAi* embryos**. Movies showing His2Av-RFP channel (orthogonal. view, left ; and maximum intensity projection, center) and MCP-GFP channel with transcription directed by *m5/m8* ^*[II]*^ (maximum intensity projection, right) in control (top), *Notch* (middle) and *α-Cat RNAi* (bottom) embryos. 0.36 μm/px XY resolution, 32×1μm slices and time resolution of 20s/frame. Anterior to the left; embryo imaged from the ventral side. Time indicates minutes from the beginning of nc14.

## Acknowledgments

We thank members of the Bray Lab for helpful discussions. Thanks to members of the Sanson lab for providing flies and advice and to Kat Millen and the Genetics Fly Facility for injections. We acknowledge the Cambridge Advanced Imaging Centre for their support, assistance in this work and use of their microscopes. This work was supported by a Wellcome Trust Investigator Award (212207/Z/18/Z) and a Medical Research Council Programme grant (MR/T014156/1) and by a PhD studentship to J.F.-S from the Wellcome Trust (109144/Z/15/Z).

## Author Contributions

J.F.-S. and S.J.B. planned the experiments; J.F.-S. conducted the experiments and analyzed the data; J.F.-S. and S.J.B. wrote the manuscript.

## Supplementary Information

**Figure S1.**
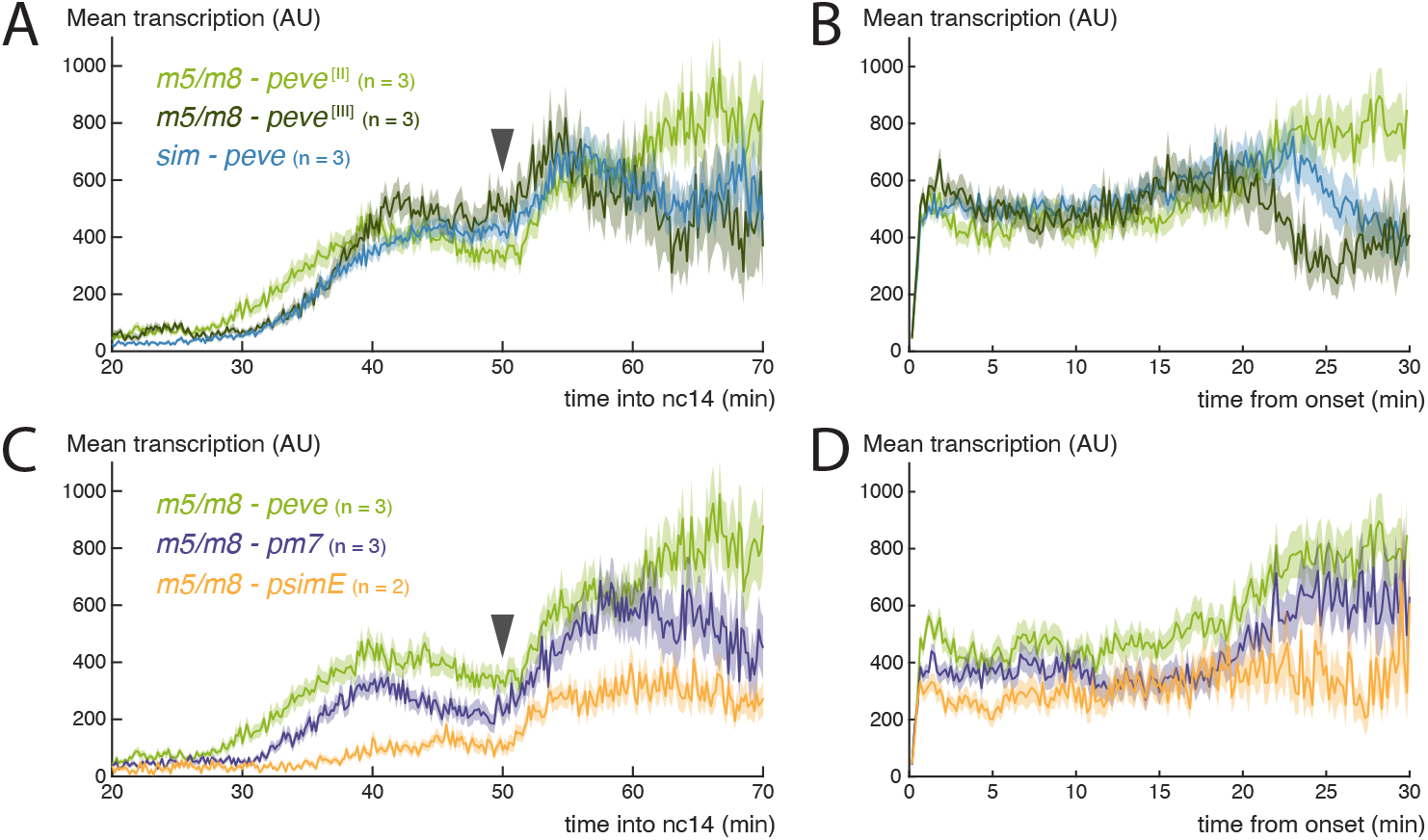
Increase in Notch dependent transcription at the time of gastrulation. **A**) Mean profile of activity of the Notch responsive enhancers *sim* and *m5/m8* inserted in landing sites in the second ([II]) and third ([III]) chromosomes. **B**) Mean profile of activity of the Notch responsive enhancers *sim* and *m5/m8* inserted in landing sites in the second ([II]) and third ([III]) chromosomes, aligned by onset times. **C**) Mean profile of activity of *m5/m8* ^*[II]*^ with different promoters (*peve, pm7* and *psimE*). **D**) Mean profile of activity of *m5/m8* ^*[II]*^ with different promoters (*peve, pm7* and *psimE*), aligned by onset times. Light green plots (*m5/m8-peve* ^*[II]*^) are the same as shown in **Fig**. 1. Mean transcription profiles show mean and SEM (shaded area) of MS2 fluorescent traces from all cells combined from multiple embryos (n embryo numbers indicated in each). Obtained by re-analizing data from ***Falo-Sanjuan et al. 2019***.

**Figure S2.**
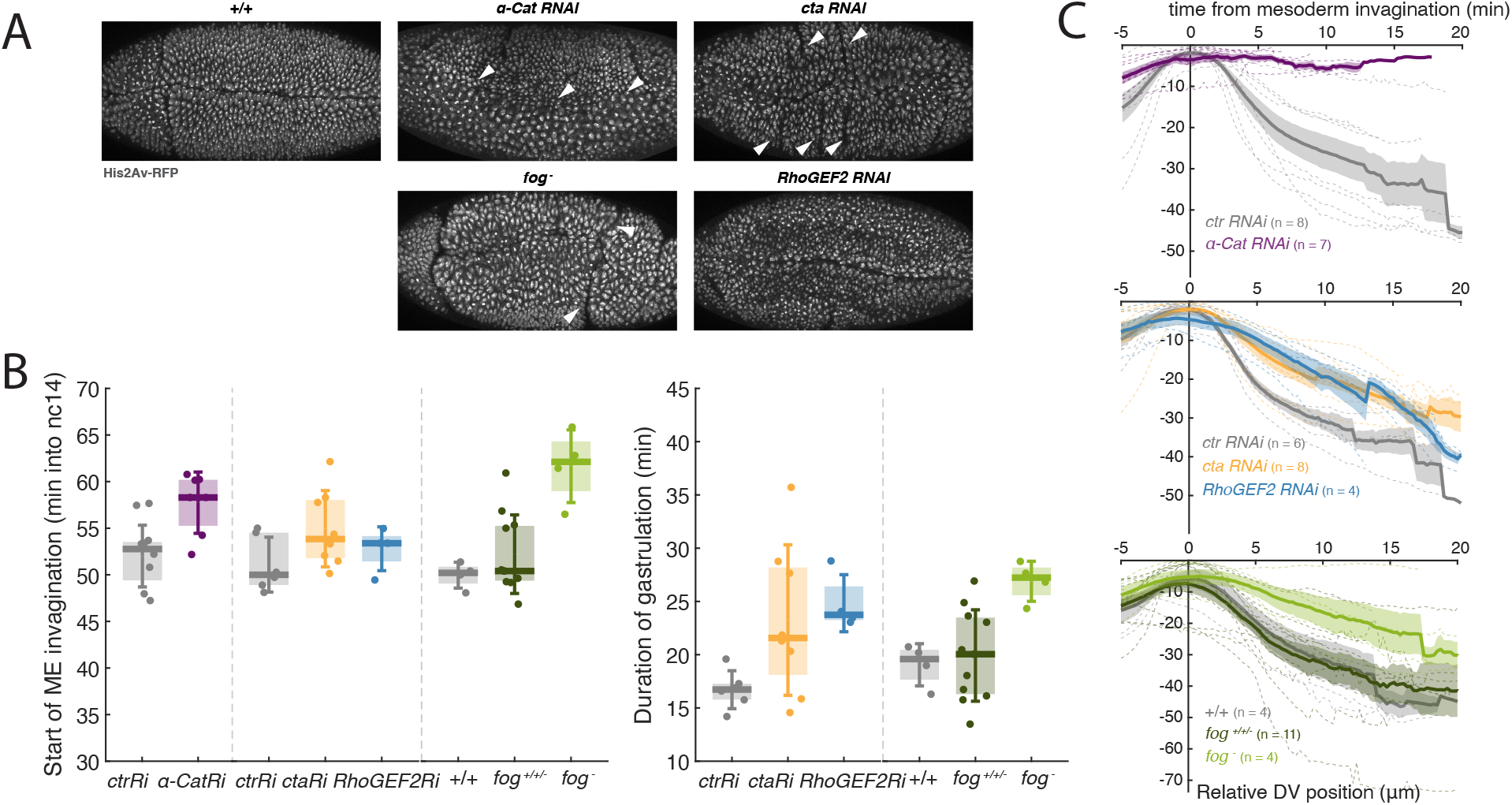
Genetic disruption of gastrulation. **A**) Still images (maximum projection of the His2Av-RFP channel) of post-gastrulation embryos from the indicated genetic conditions captured after MS2 experiments. Arrowheads indicate ectopic folds and mesoderm cells dividing externally in *α-Cat* RNAi embryos. **B**) Start of ME invagination (left) and total gastrulation length (right) in the indicated genotypes. Boxplots show median, Q1/Q3 quartiles and SD. **C**) Quantification of the movement of MSE nuclei over time in the Y axis (representing movement in the DV axis), aligned by the time of ME invagination, in the indicated genetic conditions. 0 represents the highest (most dorsal) position achieved, therefore during gastrulation MSE nuclei move towards negative positions. Mean and SEM (solid line and shaded area) of multiple embryos shown (n embryo numbers indicated in each). Dashed lines indicate mean profiles for individual embryos. Because *α-Cat* RNAi embryos do not gastrulate, total duration of gastrulation was not quantified, and mesoderm invagination has been defined, based on the overall changes occurring in the embryo, as the time when it would normally initiate.

**Figure S3.**
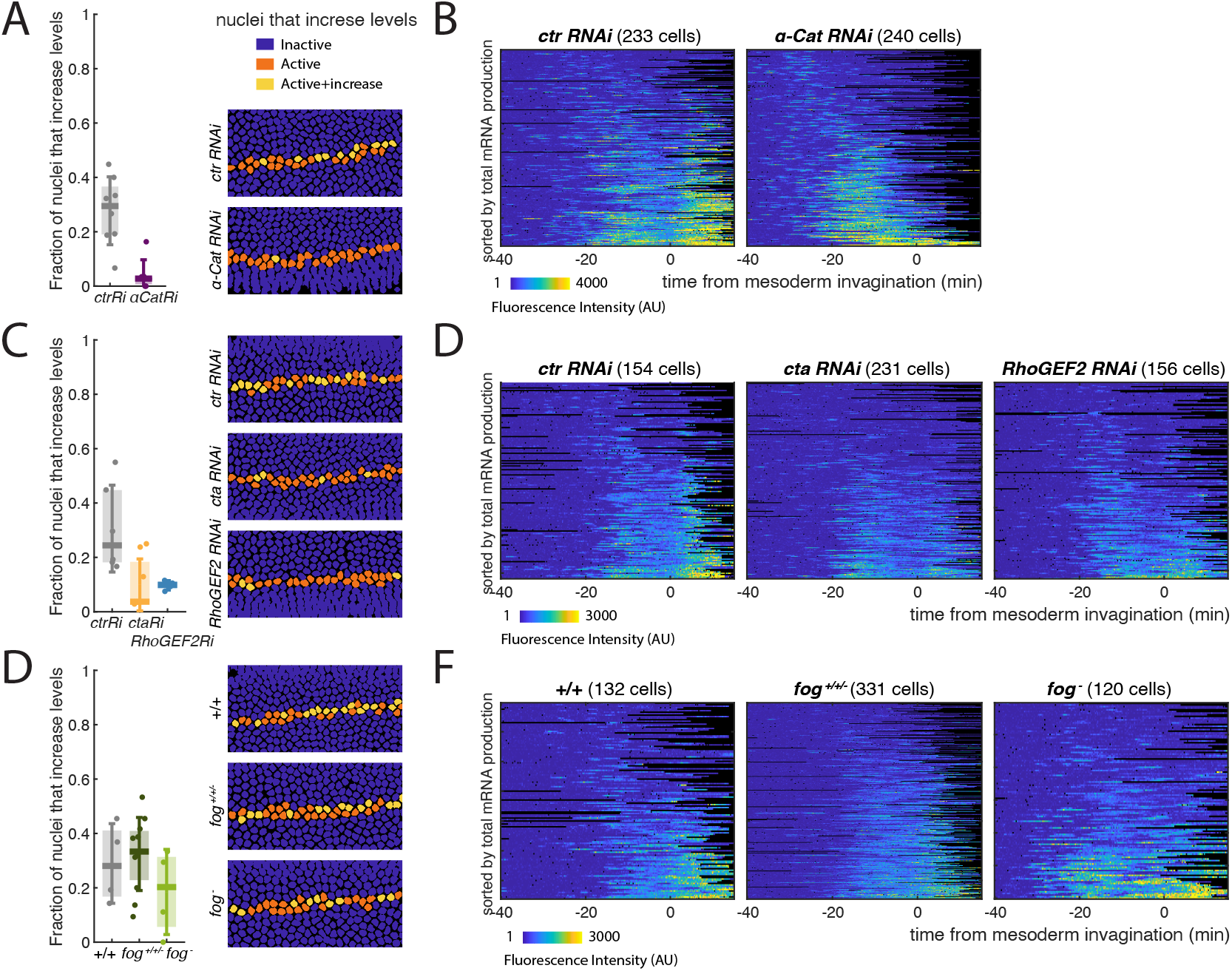
Effects of genetic manipulations to gastrulation on the increase in *m5/m8* ^*[II]*^ transcription. **ACE**) Boxplots showing proportion of active cells in each embryo that increase levels of *m5/m8* ^*[II]*^ transcription at the time of gastrulation (left; median, Q1/Q3 and SD shown) and still frame with tracked nuclei color-coded for whether they increase in levels during gastrulation (right), in the indicated genetic conditions: *α-Cat* and control RNAi (**A**), *cta, RhoGEF2* and control RNAi (**C**) and *fog*^*-*^ embryos compared to controls and other non fog hemizygous embryos obtained from the same cross - *fog*^*+/+/-*^ (**E**). **BDF**) Heatmaps showing *m5/m8* ^*[II]*^ activity in all MSE cells over time, aligned by the time of ME invagination, in the indicated genetic conditions: *α-Cat* and control RNAi (**B**), *cta, RhoGEF2* and control RNAi (**D**) and *fog*^*-*^ embryos compared to controls and other non fog hemizygous embryos obtained from the same cross - *fog*^*+/+/-*^ (**F**).

**Figure S4.**
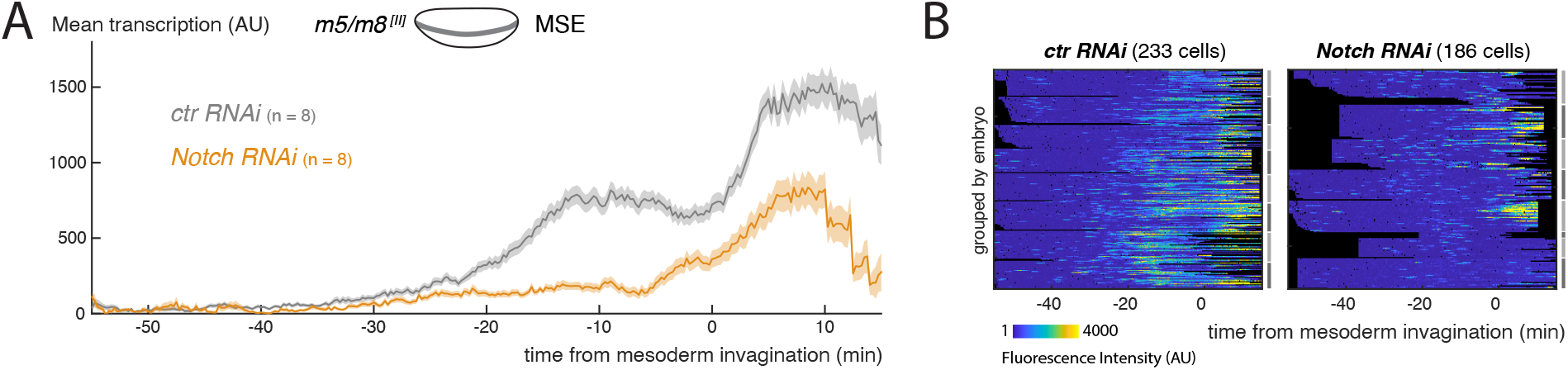
*Notch RNAi* strongly reduces Notch dependent transcription. **A**) Mean profile of transcription from *m5/m8* ^*[II]*^ in *Notch RNAi* compared to control RNAi embryos. **B**) Heatmaps showing *m5/m8* ^*[II]*^ activity cells over time in *Notch* and control RNAi embryos. Traces have been grouped by embryo (each marked by grey and black lines on the sides) to highlight the degree of embryo-to-embryo variability in transcription, likely arising from differences in the degree of Notch KD. Mean transcription profiles show mean and SEM (shaded area) of MS2 fluorescent traces from all cells combined from multiple embryos (n embryo numbers indicated in each). Control embryos are the same as shown in 2**B** and S3**B**.

**Figure S5.**
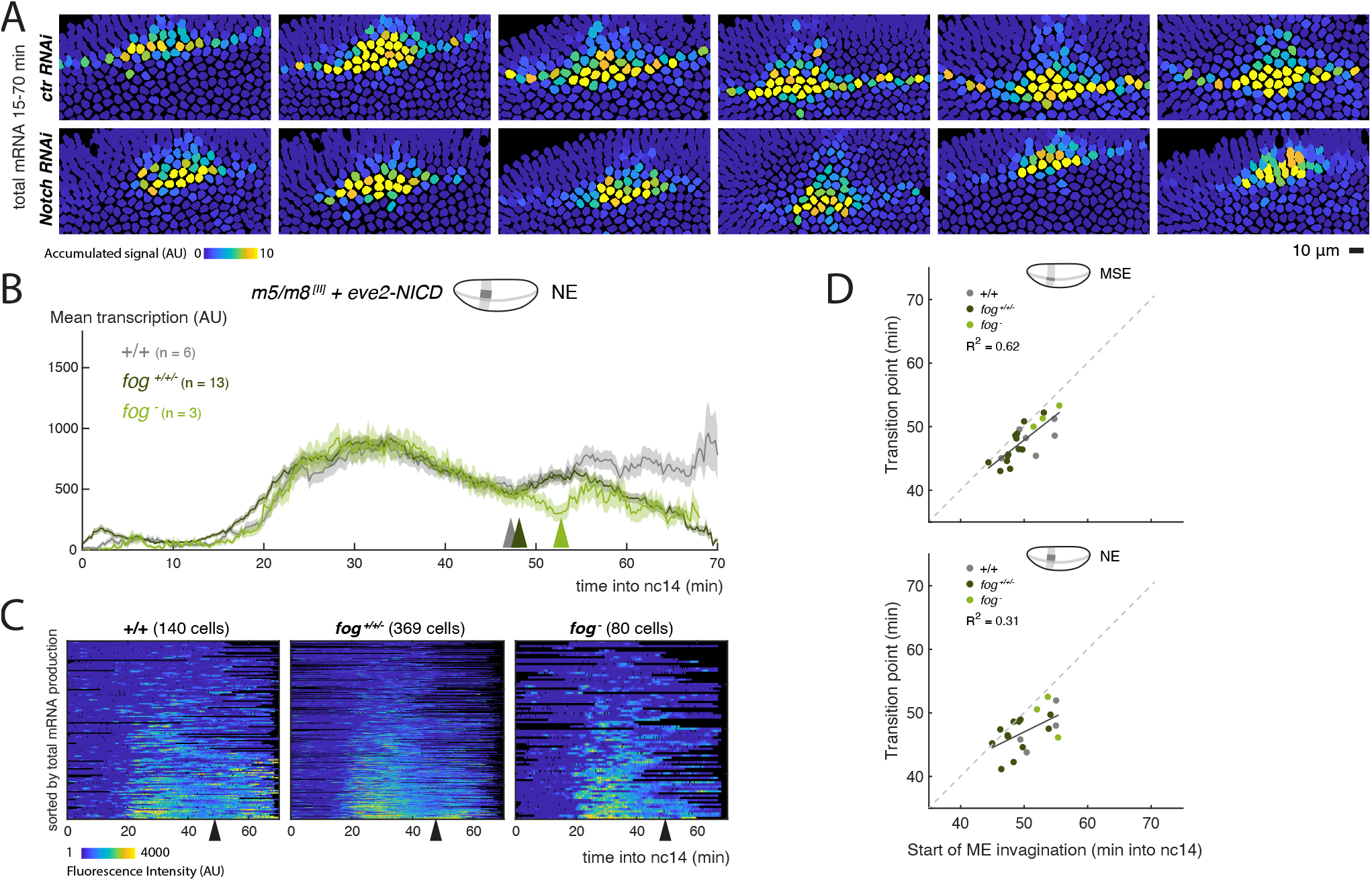
Modulation of transcription occurs downstream of pathway activation. **A**) Still frames showing tracked nuclei color-coded for the total transcription produced by each nuclei in all *m5/m8* ^*[II]*^ + *eve2-NICD* control and *Notch RNAi* embryos. Upon Notch KD, *m5/m8* ^*[II]*^ transcription only occurs where *eve2-NICD* is expressed. **B**) Mean profile of transcription from *m5/m8* ^*[II]*^ in NE nuclei expressing ectopic NICD in *fog* mutant embryos compared to controls and to other embryos obtained from the same cross (*fog*^*+/+/-*^). **C**) Heatmaps showing *m5/m8* ^*[II]*^ activity over time in all NE cells expressing ectopic NICD in *fog*^*-*^ embryos compared to controls and to other embryos obtained from the same cross (*fog*^*+/+/-*^). **D**) Correlation between start of invagination and transition in *m5/m8* ^*[II]*^ transcription levels in MSE (top) or NE (bottom) nuclei overlapping with *eve2-NICD* domain in each *fog* mutant embryo compared to control and other non *fog* hemizygous embryos obtained from the same cross. Mean transcription profiles show mean and SEM (shaded area) of MS2 fluorescent traces from all cells combined from multiple embryos (n embryo numbers indicated in each). *R*^2^ coefficients in **D** are calculated after pooling all points shown the same plot together. The transition point was only considered when a clear change in mean levels of transcription in an individual embryo could be observed, therefore the number of points could be smaller than the total number of embryos collected for each condition.

**Figure S6.**
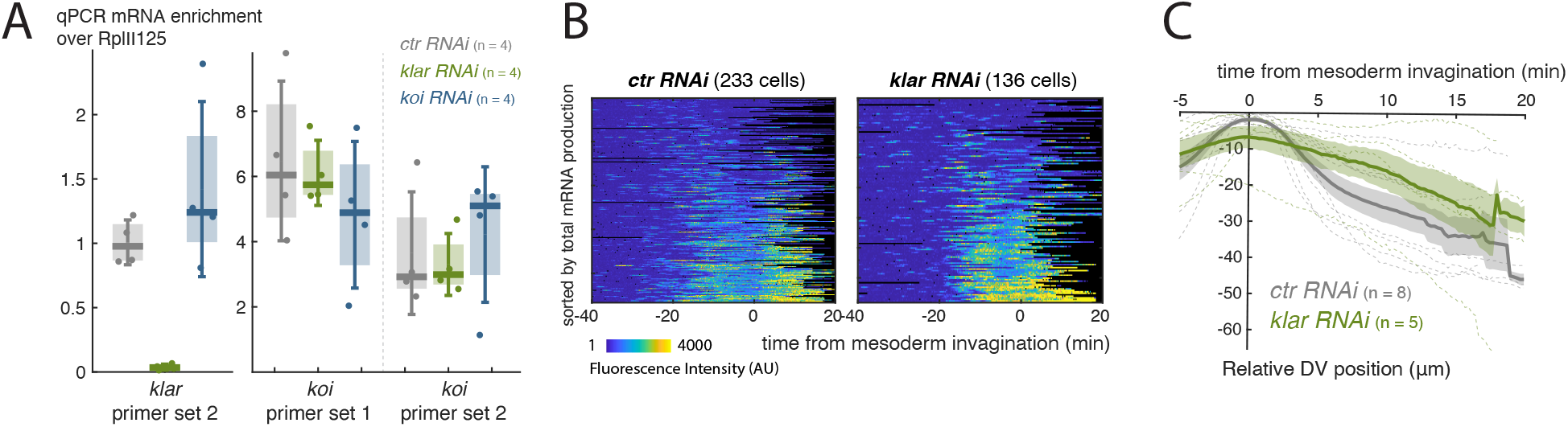
LINC complex disruption decouples gastrulation progression and changes in transcritpion. **A**) Quantification of *klar* and *koi* mRNA levels by RT-qPCR in pools of 15-20 eggs and/or pre nc13 embryos following control, *klar* or *koi* germline RNAi expression. n = 4 biological replicates for each. **B**) Heatmaps with *m5/m8* ^*[II]*^ activity, aligned by the time of ME invagination, in *klar* and control RNAi embryos. **C**) Movement of MSE nuclei over time in the Y axis (representing movement in the DV axis), aligned by the time of ME invagination, in *klar* and control RNAi embryos. Mean and SEM (solid line and shaded area) of multiple embryos shown (n embryo numbers indicated in each). Dashed lines indicate mean profiles for individual embryos. Control embryos are the same as shown in S3**B** and S2**C**.

